# Evolutionary dynamics of a quantitative trait in a finite asexual population

**DOI:** 10.1101/033597

**Authors:** Florence Débarre, Sarah P. Otto

## Abstract

In finite populations, mutation limitation and genetic drift can hinder evolutionary diversification. We consider the evolution of a quantitative trait in an asexual population whose size can vary and depends explicitly on the trait. Previous work showed that evolutionary branching is certain (“deterministic branching”) above a threshold population size, but uncertain (“stochastic branching”) below it. Using the stationary distribution of the population’s trait variance, we identify three qualitatively different sub-domains of “stochastic branching” and illustrate our results using a model of social evolution. We find that in very small populations, branching will almost never be observed; in intermediate populations, branching is intermittent, arising and disappearing over time; in larger populations, finally, branching is expected to occur and persist for substantial periods of time. Our study provides a clearer picture of the ecological conditions that facilitate the appearance and persistence of novel evolutionary lineages in the face of genetic drift.

## 1. Introduction

Speciation is said to be “ecological” when reproductive isolation has resulted from divergent natural selection driving sub-populations into different ecological niches (Schluter and Conte, 2009). When this divergence occurs in sympatry, the initial differentiation of phenotypic traits requires multiple fitness peaks in the adaptive landscape (Calsbeek et al., 2012), with selection favoring different phenotypes given the current composition of the population. Divergent natural selection, however, does not always lead to phenotypic divergence – i.e., evolutionary branching – if there is not enough variation for selection to act upon or when genetic drift is too strong relative to selection, even when the populations are asexual.

Historically, most quantitative genetic models were developed under the assumption that selection is frequency–independent with a single optimum, i.e., that fitness landscapes are constant and single-peaked. That said, the potential importance of frequency-dependent selection as a source of quantitative genetic variation was recognized early on (Clarke and O’Donald, 1964; Cockerham et al., 1972), and some quantitative genetic models have included frequency-dependent selection (e.g., Bulmer, 1980; Lande, 1976; Slatkin, 1979; Bürger and Gimelfarb, 2004). Social interactions between individuals of the same species, whether competitive, spiteful or altruistic, as well as interspecific interactions, such as interactions between predators and their preys or hosts and parasites, often result in frequency-dependent selection (Doebeli and Dieckmann, 2000). It is therefore crucial to understand how frequency–dependent selection affects the evolution of quantitative traits, under both stabilizing or diversifying selection, since the former seems to be neither more prevalent nor stronger than the latter in nature (Kingsolver et al., 2001). Particularly needed are models that incorporate both frequency-dependent selection and drift.

After the pioneering works of I. Eshel (Eshel and Feldman, 1984; Eshel, 1996), the desire to understand the long-term implications of frequency–dependence led to the development of the adaptive dynamics framework (Geritz et al., 1998; Doebeli, 2011). The method requires the assumption that mutations are rare, so that evolution proceeds as a series of competitive displacements of resident genotypes by mutant genotypes; mutations are also assumed to be of small phenotypic effect and population sizes are typically assumed to be large. Central to the framework is the concept of invasion fitness (Metz et al., 1992), which corresponds to the initial growth rate of a rare mutant in a population of very large size.

The assumption that the population size is large is a central one in the adaptive dynamics framework, but computer simulations have helped investigate the consequences of stochasticity in populations of smaller size (e.g., Dieckmann and Doebeli, 1999; van Doorn and Weissing, 2002). Because population size affects the fate of mutations, the outcome of an adaptive dynamics process can change in small populations. Claessen et al. (2007) for instance observed that evolutionary branching was much harder to obtain in individual-based simulations with small population sizes. Claessen et al. proposed two explanations for this phenomenon: first, because of random drift, the trait mean in the population changes over time and may wander away from the area where branching can happen. Second, even if branching is initiated, the incipient branches may go extinct by chance.

In populations of small size, Proulx and Day (2002) showed that the probability of fixation is a better predictor of the course of evolution in stochastic environments than invasion fitness. Similarly, the “canonical diffusion of adaptive dynamics” (Champagnat et al., 2006; Champagnat and Lambert, 2007), which describes the evolution of a quantitative trait in a finite asexual population, involved gradients of fixation probability (instead of invasion fitness). Although they allow the consideration of the effect of genetic drift, these two approaches dealt with directional selection only and did not account for the creation and maintenance of quantitative genetic diversity due to frequency-dependent selection. Obviously, as a probability of fixation refers to the fixation of one genotype and the loss of another, this measure of evolutionary success does not naturally describe the maintenance of diversity (Rousset, 2004; Allen et al., 2013). In other words, a method based on a trait substitution sequence, which assumes that the fate of a mutation is either loss or fixation, is not suited to account for evolutionary diversification, where different types coexist.

In this article, we study the evolution of a quantitative trait under frequency–dependent selection, in an asexual population of finite, but not fixed, size. We use a moment-based approach, because it bridges the gap between quantitative genetic and adaptive dynamic frameworks (Abrams et al., 1993; Abrams, 2001; Débarre et al., 2013; Débarre et al., 2014). We illustrate our results with a model of social evolution in a well-mixed population (i.e., in the absence of any spatial or social structure), where the quantitative trait *Z* under selection corresponds to investment in social behavior (Doebeli et al., 2004; Lehmann, 2012; Wakano and Lehmann, 2012; Wakano and Iwasa, 2013).

Our study builds upon the work of Wakano and Iwasa (2013). In their model, Wakano and Iwasa (2013) assume asexual reproduction, discrete, non-overlapping generations and a potentially small but constant population size (using a Wright-Fisher model). The authors explore models where branching is expected in infinite populations but may fail to occur within finite populations. They identify two major parameter regimes involving diversifying selection: where branching is expected deterministically and continues to be observed in finite populations even if mutations have small effects (termed “deterministic branching”) and where branching is expected deterministically but will only occur in finite populations occasionally, when mutations are of large enough size to overwhelm drift (termed “stochastic branching”).

Here, we extend the framework of Wakano and Iwasa (2013) to populations whose size is finite but not fixed and to a lifecycle with overlapping generations (a birth-death process). We derive expressions for the stationary distribution of the total population size, trait mean, and trait variance under stabilizing selection, and we show how these distributions can help us refine the conditions for evolutionary diversification when selection is diversifying. In particular, we show that the “stochastic branching” regime identified by Wakano and Iwasa can be sub-divided further into (i) a “no branching” regime in which branching will either never occur or be so seldom and collapse so rapidly that the population is very unlikely to be observed in a diversified state; (ii) an “intermittent branching” regime in which branching arises and collapses over biologically reasonable time frames; and finally (iii) a regime akin to the “deterministic branching” regime, in which branching is so likely and collapses so rarely that the system maintains multiple species almost always, with populations likely to remain branched for long enough to accumulate further speciation barriers.

## 2. Model and methods

### 2.1. Model

We describe the evolution of a trait *Z* in a population of asexual individuals. Each individual in the population is characterized by its genotype *z_i_*, which we also refer to as phenotype in the absence of environmental effects; in the remainder of the article, we refer to *z_i_* as “type” or simply “trait”. At a given time *t*, we denote the current size of the population by *N*(*t*), while a vector **z**(*t*) summarizes all the types present in the population. The trait mean (first moment of the distribution) is 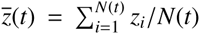 and the variance (second central moment of the distribution) is 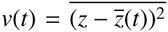. For each of these variables, we may drop the time dependency for simplicity. Each time step, either one individual reproduces (producing exactly one offspring) or one individual dies.

We use the term “fecundity” to refer to the reproductive potential of an individual, which is proportional to the chance that this individual will reproduce in a time step. We assume that individual fecundity *F_i_* depends on both the type of each individual and the distribution of types in the population; we denote by 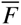 the mean fecundity in the population. When it does reproduce, a parent of type *z_i_* produces an offspring with phenotype *z_i_ + δ*, where *δ* follows a distribution *u* (called a mutation kernel) with mean *m_u_ =* 0 and variance 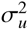 assumed to be small. Mutation is therefore a source of variation in the population, and multiple types can coexist at any time point even though the population is finite and potentially small.

Individual survival, on the other hand, is type– and frequency– independent, but it is density–dependent: individual survival decreases as the size of the population increases. The *per capita* death rate per time step, *D_i_*, is defined as *D_i_ = dN*. We denote by 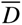 the mean death rate in the population.

In the next time step, both the size of the population and the distribution of types have changed; our aim is to find expressions for their stationary distributions. Key to our derivation is the assumption that populations that have not diversified have a trait distribution that is Gaussian (at any time), with a small variance v. Hence, we only need to follow the mean 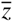 and variance *v* of the distribution of types. By contrast, populations undergoing evolutionary branching are characterized by a substantial increase of trait variance. Thus, we determine whether or not evolutionary branching is likely to be observed by determining when the steady state distribution for the trait variance does not or does have substantial density at small values of *v*.

#### Illustration: social evolution

We illustrate our results using the specific example where *Z* is a social trait that represents individual investment into social behavior; the trait can take any value between 0 (no investment) and 1 (maximum investment). Initially analyzed by Doebeli et al. (2004) under the assumption that population size was infinite, this model has also been used by related studies (Wakano and Lehmann, 2012; Wakano and Iwasa, 2013), in which population size, now finite, was fixed. Although widely used in population genetics, a fixed population size is a restrictive assumption; in particular, it requires that all the deaths that may occur within a time step are exactly compensated by the same number of births (e.g., *N* deaths and *N* births in a Wright-Fisher model; one death and one birth in a Moran process). In our model, we do not impose a fixed population size. Because either a death or a birth happens within one time step, population size does change with time. We will see, however, that the dynamics of the population size occur on a much faster time scale than the dynamics of the mean and variance of the distribution of traits; population size hence reaches a quasi-equilibrium 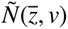, and can therefore be approximated as being stabilized when changes in 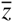 and *v* are considered. For the sake of comparison, we use in our study the same payoff values as in the previous studies that our work complements.

The population is well-mixed: each individual interacts socially with every individual in the population (i.e., “playing against the field”), which includes the influence that a focal player itself has on that field; these social interactions affect their fecundities. Using payoff functions similar to the ones in Doebeli et al. (2004), we can rewrite the fecundity of an individual with trait *z* as follows:

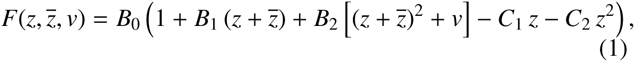

where 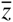 is the current mean type and *v* is the current variance. The parameters *B*_1_ and *B*_2_ (resp. *C*_1_ and *C*_2_) refer to benefits (resp. costs) of social interactions, while *B*_0_ is the baseline fecundity. The steady state population size can thus be altered by varying *B*_0_, while keeping the relative strength of frequency-dependent interactions constant.

### 2.2. Method

#### 2.2.1. Diffusion approximations

We want to find the stationary distributions of three quantities: the size of the population (*N*), the trait mean 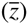 and variance (*v*). To do so, we use diffusion approximations. Diffusion approximations rarely yield results for multiple variable models, but it turns out that changes in the three variables occur at different time scales, so that we can decouple them and derive diffusion approximations for each variable separately.

To derive the diffusion approximation for a variable, we need expressions for the expected change in that variable between two time steps (which we denote by the letter *μ*) and also the expected squared change (which we denote by σ^2^). Once we have derived these terms, the stationary distribution of the variable of interest is given by

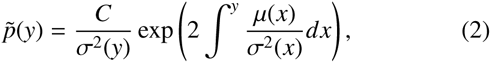

the constant *C* being chosen such that 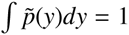 (since 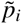 is a probability density) (Rice, 2004; Otto and Day, 2007).

Key to the derivation is the assumption that the variance in trait values, *v*, is small, implying that the distance between the trait of any individual and the trait mean is small. As a result, we can expand individual fecundity *F* as follows:

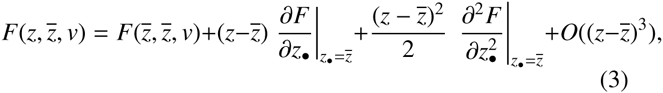

where *z_●_* refers to the first argument of *F*. Similar expressions are commonly used in the inclusive fitness framework; many of these studies focus solely on the first order term and hence cannot be used to determine whether evolutionary branching will occur (as noted by Lehmann and Keller, 2006, see references therein for extensions within the inclusive fitness framework that investigate branching and do include second order terms). For notational shorthand, we define:

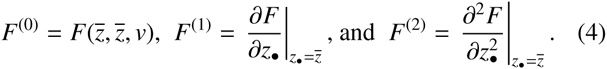

We note that in the example of social evolution, individual fecundity *F* is already a quadratic function of *z* (see equation (1)), so that neglecting 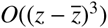 in equation (3) will not constitute an approximation.

#### 2.2.2. Stochastic simulations

The stationary distributions are then compared with the results of individual-based stochastic simulations of the model. The simulations are written in C and analyzed in R (R Development Core Team, 2011). The trait space is divided into 51 discrete values between 0 and 1, so that the phenotypic distance between two adjacent trait values is *dz* = 0.02. Mutation occurs with probability *μ*_0_ = 0.01 and changes the trait value into the value of one of the two adjacent phenotypes (*z′ = z* ± *dz*). With this mutation kernel, the mutational variance is 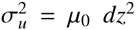 (except at the edges of the trait space), and we use the same parameter values as in Wakano and Iwasa (2013).

The simulation codes have been uploaded on the data repository Dryad (Provisional DOI: doi:10.5061/dryad.b3604).

## 3. Results

### 3.1. Deriving the terms of the diffusion approximations: Expected changes and squared changes

In the following, we calculate the moments (*μ* and σ) needed to determine the stationary distribution for the population size *N*, the trait mean 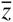, and the trait variance *v*. Because we are primarily interested in the stationary distribution and not the temporal dynamics, we consider the process only at time points at which an event occurs (either a birth or a death): a “time step” corresponds to the occurrence of one such event, and we are now focusing on changes in population size, trait mean, and trait variance occurring during a time step. More detailed calculations of all of these terms are presented in Appendix A.

#### 3.1.1. Changes in the population size

The change in population size is +1 if one individual reproduces and −1 otherwise; the expected change in population size is

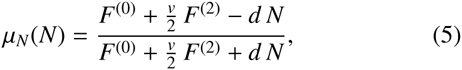

and the expected squared change in population size is

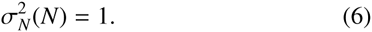

(See Appendix A for details of the derivation.) Given appropriate values for 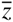 and *v*, equations (5) and (6) can be used in (2) to describe the stationary distribution for the population size.

While the changes in population size are of order *O*(1), we shall see in the next two sub-sections that changes in the distribution of traits are of order *O*(*v*) or higher (equations (8) and (11)). We can therefore consider that population size reaches and hovers around a mean quasi-equilibrium value 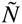 before further changes in *z* and *v* occur; 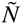 is defined such that 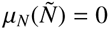:

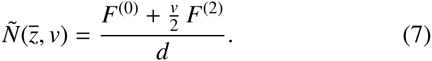

In the remainder, we will hence assume that the population is at this demographic equilibrium. This implies, in particular, that the chance that the next event is a death is equal to the chance that the next event is a birth; or, in mathematical terms, that 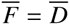.

#### 3.1.2. Changes in trait mean

The expected change in the trait mean is

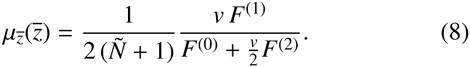

(See Appendix A for details of the derivation.) Because mortality is type–independent, death events do not change the trait mean, on average. Birth events do and happen one-half of the time at demographic equilibrium, but only the newly born individual affects the trait mean when birth occurs, hence the 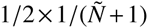 factor in (8). This expected change in trait mean is of order *v*, the current variance in the population, which is assumed to be small; this justifies the separation of time scales between changes in the trait mean and changes in population size (equation (5)). The remaining factor in (8), is a selection gradient, the denominator representing the current average fecundity in the population. A Price equation (Price, 1970) version of (8) is presented in the Appendix, equation (A.16b).

The expected squared change in the trait mean is given by

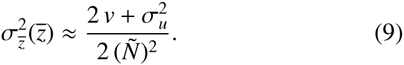

(Details of the calculations are presented in Appendix A.) Given appropriate values for 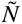 and *v*, equations (8) and (9) can be used in (2) to describe the stationary distribution for the trait mean.

The expected change in the trait mean (equation (8)) is zero when *vF*^(1)^ = 0, that is, either when there is no variance in the population—which we ignore because mutation always regenerates variance—or when the selection gradient *F*^(1)^ vanishes (see (4)). In our example, with the fecundity function defined as in (3), this occurs when the trait mean is at 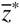,

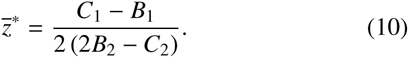

#### 3.1.3. Changes in variance

We now consider changes in the variance of the distribution of traits in the population at demographic equilibrium. Using the fact that the distribution is Gaussian, we eventually obtain the following expected change in the trait variance:

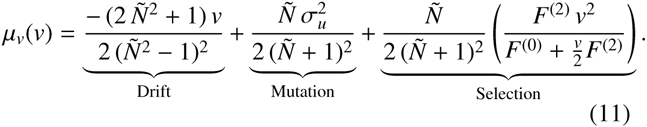

(See Appendix A for details of the calculations.) At demographic equilibrium, birth and death each occurs with a chance 1/2, hence the overall 1/2 factor in equation (11). The first term in equation (11) corresponds to the effect of random drift, which reduces the variance, and is stronger in smaller populations. In a large enough population, the effect of drift on the trait variance is of order 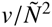, which is negligible compared to the effect of selection on the trait mean (equation (8)), justifying a separation of time scales. This may however not be the case in very small populations. The second term in equation (11) corresponds to the effect of mutation, via the variance of the mutation kernel, 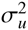. The effect of mutation on the trait variance is of order 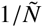 because there is at most one new mutant between the time steps. Finally, the third and last term in equation (11) corresponds to the effect of selection. Its sign is the sign of *F*^(2)^, the curvature of the function *F*. Selection is stabilizing when *F*^(2)^ < 0 and tends to reduce the trait variance in the population. Selection is diversifying, i.e., tends to increase the trait variance, when *F*^(2)^ > 0. A Price equation (Price, 1970) version of (11) is presented in the Appendix, equation (A.23).

For the expected squared change in the trait variance, still assuming a Gaussian distribution of traits with a small variance and a relatively large population size, we obtain

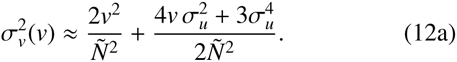

(Details of the calculations are presented in Appendix A.) Numerical comparisons (in the *Mathematica* file provided as supplementary material on Dryad^1^) show that the shape of the stationary distribution of *v* is little affected if we neglect the effects of mutation on the squared change in trait variance and use the simpler equation

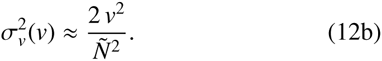

Again, given appropriate values for 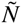 and 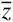, equations (11) and (12b) can be used in (2) to describe the stationary distribution for the trait variance.

In the absence of selection, the last term of equation (11) disappears. We denote by 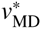 the variance at the mutation-drift equilibrium, i.e., the variance such that *μ_v_*(*v*_MD_) = 0 when *F*^(2)^ = 0.

### 3.2. Stationary distributions under stabilizing selection

The final equilibrium values of the population size, trait mean, and trait variance, 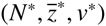, are obtained by solving the system 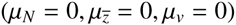.

#### 3.2.1. Population size

The stationary distribution of the population size is obtained by integrating equation (2) with the diffusion terms *μ_N_* and 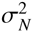, setting 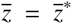 and *v* = *v*^*^. Under stabilizing selection, the equilibrium variance is very small, and similar distributions of *N* are obtained if we set *v* = 0 instead of *v*^*^; we will therefore hereafter approximate the equilibrium population size by 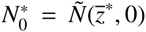. The solution is compared to the output of individual-based simulations in figure 1(a) and (d).

**Figure 1:**
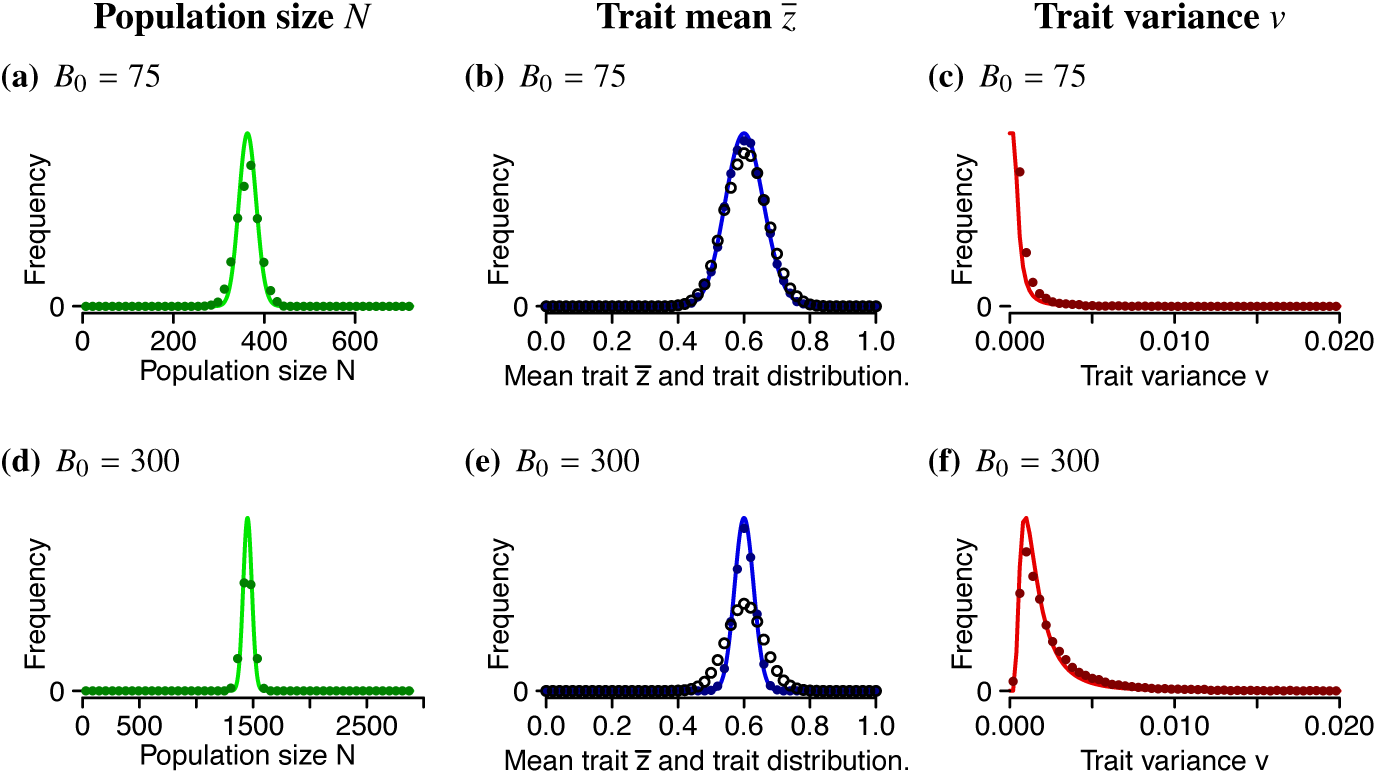
Stationary distributions of the three variables of interest when selection is stabilizing. In the first row, *B*_0_ = 75, in the second row, *B*_0_ = 300. Other parameters in all figures: *B*_2_ = −1.5, *B*_1_ = 7, *C*_2_ = −1, *C*_1_ = 4.6, *d* = 1; *μ*_0_ = 0.01 and *dz* = 0.02, so that *σ_v_* = 0.002. The curves are the analytic predictions; solid colored dots are simulation results for comparison, while black open dots in panels (b) and (e) show the entire trait distribution itself (i.e., not the distribution of trait means)—both are averaged across time for one simulation.

#### 3.2.2. Trait mean

The stationary distribution of the trait mean is obtained by integrating equation (2) with the diffusion terms 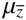 and 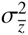, setting 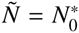 and *v* = *v^*^*. The solution, together with the output of individual-based simulations, is presented in figure 1(b) and (e). In each case, the expected distribution of the trait mean estimated from the simulations (solid dots) and the analytical prediction (solid curve) match well. Also shown in these figures is the distribution of trait values (open dots), which is wider than the distribution of trait means, especially in larger populations (panel (e)), because of the variance maintained within populations.

In smaller populations, the steady state distribution for the trait mean is more broadly distributed and makes more excursions away from the optimum at 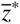 (as in Claessen et al., 2007). This can be seen by comparing figure 1(b), which has a lower population size due to a lower baseline fecundity *B*_0_, to figure 1(e), which has a higher baseline fecundity (in both cases 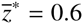).

#### 3.2.3. Trait variance

The stationary distribution of the trait variance is obtained by integrating equation (2) with the diffusion terms *μ_v_* and 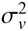, setting 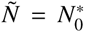 and 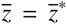. This solution is compared to the output of individual-based simulations in figure 1(c) and (f). More variance is maintained in larger populations (panel 1(f) with a higher baseline fecundity *B*_0_). By contrast, drift depletes the trait variance in smaller populations (panel 1(c)).

### 3.3. Does diversification occur, if selection is diversifying?

The results that we have derived so far rely on the assumption that the distribution of traits is approximately Gaussian with small variance. These assumptions may not hold when selection is diversifying (i.e., when *F*^(2)^ evaluated at 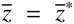 is positive), in which case evolutionary branching may occur, causing the distribution of traits to become bimodal. Whether this branching actually occurs, however, depends on the extent of random drift. In the remainder, we will use our results on the expected change in the variance and the stationary distribution of the trait variance, both derived assuming a Gaussian distribution of traits with small variance, to identify conditions that violate the assumption that variance remains small and hence imply that evolutionary branching does occur, despite drift.

The rationale is that until evolutionary branching—if it ever happens—the equations that we derived previously are still valid. We assume that at the time at which branching happens (if ever), the population size is *N*^*^ and the trait mean is 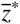. We then evaluate the steady state variance, relative to that expected at mutation-drift equilibrium, 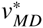 (i.e., the equilibrium value of the variance in the absence of selection (see equation (A.26))). We then classify branching regimes in two steps. We first identify when the mean of the variance approaches a small value (branching not possible) and when it can increase to high values (branching possible). In the latter case, we then approximate the steady state distribution for the variance to determine the proportion of time that the population spends in the low variance (unbranched) state and in the high variance state (branched).

Note also that we are considering the variance of the distribution of traits in the population, but not the actual shape of this distribution. That is, we only determine the conditions under which the variance becomes large, but we cannot prove that the resulting distribution is bimodal. The deterministic version of the model of social evolution does show, however, that diversifying selection leads to branching, i.e., to the evolution of distinct clusters of traits (see Appendix D). We will therefore still write about branching when the variance grows substantially.

#### 3.3.1. Using the expected change in the variance *μ_v_*

We next determine the conditions under which the trait variance will increase when selection is diversifying; the answer is given by the sign of *μ_v_*, which depends on the total population size and hence on the baseline fecundity *B*_0_. As (11) is quadratic in *v*, *μ_v_*(*v*) = 0 admits from zero to two admissible solutions depending on the value of *B*_0_; numerical examples are given below using the parameters of figure 2.

**Figure 2:**
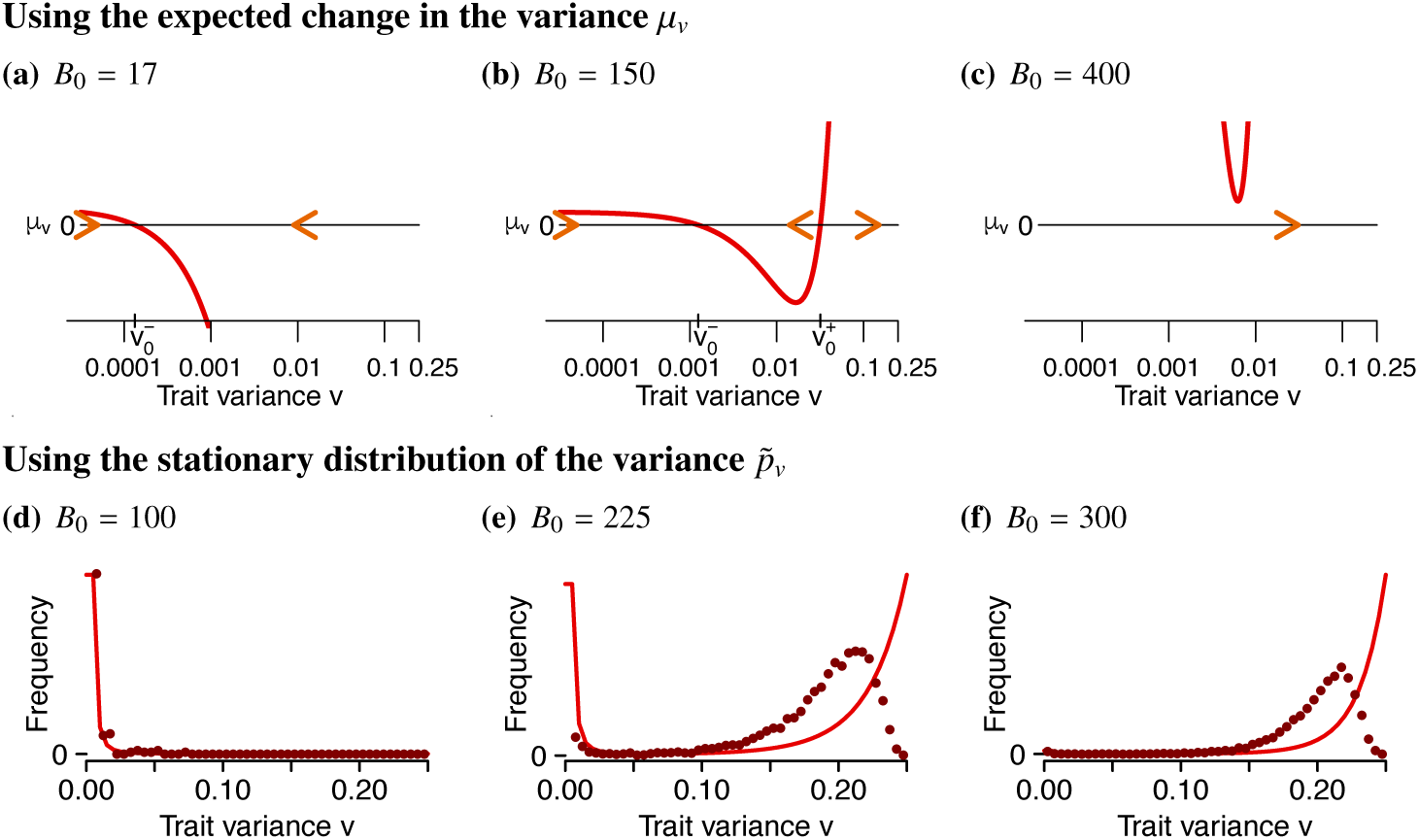
The effect of baseline fecundity *B*_0_ on branching when selection is diversifying. (a)–(c): Expected change in the trait variance, *μ_v_*, as a function of the current variance within the population, *v* (on a log scale). (d)–(f): Stationary distribution of the trait variance, 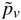 (curve: analytical predictions derived under the assumption of a Gaussian distribution of traits of small variance; dots: output of individual-based simulations).

For low values of *B*_0_ (*B*_0_ < *B_μ_*,*_L_*; *B_μ_,_L_* ≈ 19 for the parameters used in figure 2), *μ_v_*(*v*) admits only one root, 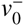, in the interval of possible values of *v* for a trait bounded between 0 and 1 (0 ≤ *v ≤* 0.25), and this equilibrium is stable (see figure 2(a)). The variance 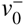 is slightly greater than 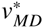, the variance at mutation-drift equilibrium. This means that the equilibrium trait variance will remain very low: the small population size leads to strong random drift, which counteracts the effects of diversifying selection. In this case, branching will not be observed despite being selectively favored.

For intermediate values of *B*_0_ (*B_μ_,_L_ ≤ B_0_ ≤ B_μ_*,*_H_*; for the parameters considered, *B_μ_*,*_H_* ≈ 394), *μ_v_* admits two roots, 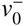 and 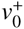, and *μ_v_* is positive for 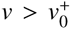; hence, 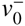 is locally stable, and 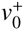 is unstable. If the current trait variance within the population is very low, then the variance will equilibrate at the low value 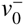. If, however, the trait variance makes an excursion and reaches a high value (above 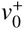), e.g., due to the appearance of mutations with larger than average effects or simply due to drift, then the variance will keep increasing (see figure 2(b)) and branching will result.

For higher values of *B*_0_ (*B*_0_ > *B_μ_*,*_H_*), *μ_v_* is always positive. Consequently, the variance is always expected to increase regardless of its current value, and diversification is certain (see figure 2(c)).

This latter regime was called “deterministic branching” by Wakano and Iwasa (2013), while the regime they called “stochastic branching” corresponds, in our model, to both of our first two regimes (with B_0_ < *B_μ_,_L_* and *B_μ_,_L_ ≤ B_0_ ≤ B_μ_,_H_*). Here we have shown that if there are limits to the maximum amount of variation that can be introduced by mutations (e.g., because the trait is bounded), then populations that are too small (*B*_0_ < *B_μ_,_L_*) are never expected to branch even though selection is diversifying (*F*^(2)^ > 0).

We are now going to refine the description of what happens for the intermediate values of the baseline fecundity *B*_0_, that is, for intermediate population sizes.

#### 3.3.2. Using the stationary distribution of the variance

While the inspection of *μ_v_* allowed us to determine when stochastic branching is plausible, how often the branches arise and how quickly they collapse remain unclear. In this section, we use, as an approximation, the stationary distribution of the variance (see figure 2(d)–(f)), to ask how often the variance is likely to be small (near 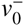) and how often large (near 0.25), corresponding to unbranched and branched states of the population, respectively. We must, however, keep in mind that the stationary distribution is derived under the assumption of a Gaussian distribution of traits with small variance, which will not hold after branching occurs because the population becomes bimodal. Nevertheless, the Gaussian approximation primarily causes errors in estimating the amount of variance when in the branched state (see e.g., figure 2(f)), while it works well to determine how often the system is likely to be in the state with small variance (see figure 2(d)).

For extremely low values of *B*_0_ (*B*_0_ < *B_μ_,_L_*), inspection of *μ_v_* showed that branching cannot occur; when *B*_0_ remains low but is above *B_μ_,_L_*, we saw previously that the variance could grow if it ever made an excursion to a high enough value (once 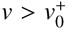 so that *μ_v_* > 0), but the stationary distribution of *v* indicates that these transitions remain extremely rare and short-lived (see figure 2(d)).

For intermediate values of *B*_0_, the stationary distribution of the trait variance is itself bimodal (see figure 2(e)): the variance may reach (or oscillate between) different values: a low value corresponding to a unimodal distribution of the trait (no branching) and a high value corresponding to a bimodal trait distribution (after branching).

For higher values of *B*_0_, the stationary distribution indicates that the trait variance is mostly, or even exclusively, high: branching will happen and tend to persist (see figure 2(f)). This occurs when *B*_0_ is high enough, even if *B*_0_ < *B_μ_,_H_*, so that the variance could decline (*μ_v_* < 0) if ever the variance made an excursion to a low value 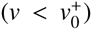. We see from the stationary distribution of *v* that these transitions remain extremely rare when *B*_0_ is large enough: the variance will almost always be high.

To better characterize the transitions between these three regimes as a function of the population size (altered by adjusting the baseline fecundity *B*_0_), we plot the probability that the variance is greater than 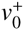, the unstable equilibrium variance identified in the previous section using the expected change in the variance *μ_v_*. This probability is determined using the stationary distribution of the variance: denoting by *p_v_* the stationary distribution of the variance, we compute 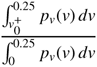 for different values of *B*_0_. The result is plotted in figure 3 (curve), and compared to the outcome of stochastic simulations (dots). Here again, the exact values of the boundaries between these regimes depend on the model and parameter values. From Figure 3, we can clearly identify the three regimes: for low values of the baseline fecundity (*B*_0_ < *B_p,L_;* 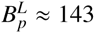 with the parameters of figure 2), the variance in the population remains mostly around its low value 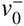 (it is only 1% of the time above 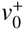), and drift nearly always overwhelms diversifying selection. For intermediate values of *B*_0_ (*B_p L_* ≤ *B*_0_ ≤ *B_p_,_H_*; with these parameters, *B_p,H_* ≈ 319), populations will transition back-and-forth between unbranched and branched states, which we refer to as “intermittent branching”. Finally, for sufficiently high values of *B*_0_ (*B*_0_ > *B_p_,_H_*), selection nearly always drives diversification, with the variance above 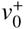 more than 99% of the time. While the precise transition points are slightly shifted in the simulations relative to the analytical predictions, the correspondence is good given the pretty restrictive assumptions (Gaussian, small variance) used to calculate the steady state distributions.

**Figure 3:**
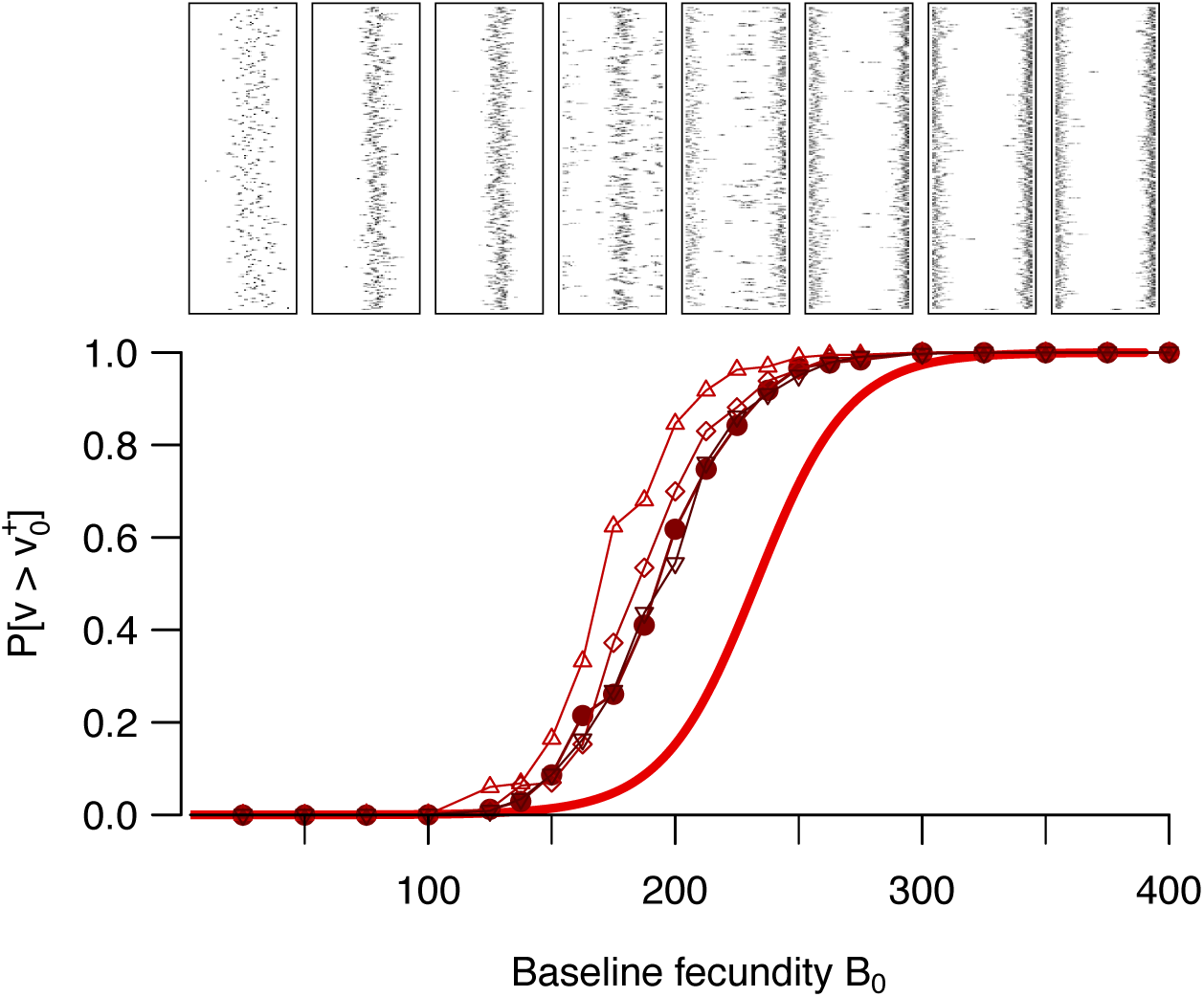
Probability that the variance is greater than 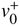, calculated using the stationary distribution of the variance 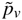 (curve) and in the stochastic simulations (dots). In mathematical terms, the curve represents 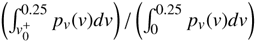. The insets above the graph represent the distribution of traits over time (horizontal axis: trait value [0 ≤ *z* ≤ 1], vertical axis: time); the horizontal location of each inset refers to the corresponding value of *B*_0_. We also explored the impact of binning the traits into different numbers of bins through simulation: 51 bins as in the remainder of the paper (circles); 11 bins (upward triangles); 26 bins (diamonds); 101 bins (downward triangles). Parameters: same as in figure 2.

In our simulations, we used parameter values that have been used in previous studies (Doebeli et al., 2004; Wakano and Lehmann, 2012; Wakano and Iwasa, 2013), which correspond to weakly diversifying selection (low *F*^(2)^) and a small variance of the mutation kernel (low *σ_μ_*). The boundaries between the three regimes identified above also depend on the choices made for these parameters; in figure S1 we show that both stronger diversifying selection (figure S1(b)) and higher mutational variance (figure S1(a)) shift the boundaries towards lower population sizes (lower *B*_0_ values), increasing the parameter range in which diversifying selection overwhelms drift.

## 4. Discussion

Using a moment-based approach and diffusion approximations, we have derived expressions for the stationary distributions of population size, trait mean, and trait variance for a quantitative trait in a well-mixed population, illustrating our results with a model of social evolution. The population was updated following a birth and death process: during a time step, either exactly one individual dies or one individual reproduces, producing one offspring. The stationary distributions were derived under the assumption that the trait distribution is Gaussian, with little trait variance, and a finite but not too small population. Assuming a Gaussian distribution allowed us to focus on the mean and variance of the distributions, all other moments being then known. The other assumptions allowed for a decomposition of time scales: population size first reaches a quasi-equilibrium, then the trait mean equilibrates, finally the trait variance reaches a steady state. We were therefore able to derive diffusion approximations for the three variables and checked them against the output of individual-based simulations, under stabilizing selection. We then used these results to derive conditions for evolutionary branching, when selection is diversifying—a regime under which the assumption of Gaussian distributions should eventually break down. Nevertheless, we find good agreement for the conditions under which branching is likely to occur, assuming that the initial unbranched population is nearly Gaussian.

A few studies (Lehmann, 2012; Wakano and Lehmann, 2012; Wakano and Iwasa, 2013) have previously investigated a similar question, using either different approaches or different assumptions.

Lehmann (2012) incorporated genetic drift into a similar model of quantitative trait evolution. The focus of this study was to determine the stationary distribution for the trait value in a single population, using a separation of time scales approach that described successive allele replacements, from one monomorphic state to another. This approach describes how often a species would be found near an evolutionary attractor and how strong convergence to this attractor would be in a finite population. The study, however, did not attempt to look at evolutionary branching.

Wakano and Lehmann (2012) extended this work, exploring the dynamics of the system assuming that a maximum of two alleles would be present at any one point in time (requiring a very small mutation rate). To assess whether invasion of a mutant allele was successful or not, the authors considered the average allele frequency across the stationary distribution, *ρ*_A_, comparing it to 1/2, its value when the mutation is neutral. Wakano and Lehmann found that, in their two-allele model and considering only very small step mutations in a finite population, the conditions for evolutionary stability were the same as the conditions for convergence stability, suggesting that branching would not occur. As shown in Appendix C, the conditions for evolutionary stability are always the same as the conditions for convergence stability using *ρ_A_*, regardless of population size or the exact life cycle. Furthermore, the authors showed through simulations that their two-allele criterion succeeded only in predicting a lack of branching at very small population sizes; it did not identify the branching that occurred at moderate population sizes (*N* = 1000), where many more than two alleles tended to segregate. Thus, this work left open the question of how to predict branching in finite populations.

The most recent of these studies, by Wakano and Iwasa (2013), used a different, moment-based approach. The authors investigated a Wright-Fisher model, with discrete, non-overlapping generations and a fixed population size (although we did not impose a fixed population size, our results indicate that population size rapidly reaches a quasi-equilibrium). Defining evolutionary branching as a situation in which the trait variance “explodes”, they were able to distinguish between regimes of diversifying selection under which branching is certain (“deterministic branching”) or not (“stochastic branching”). Using the stationary distribution of the variance, we were able to refine the conditions under which diversification can occur. In particular, we show that that “stochastic branching” regime of Wakano and Iwasa (2013) actually includes three regimes: a “no branching”, an “intermittent branching”, and a “branched” regime. In the first and last of these regimes, the population is nearly always unimodal (“no branching”) or nearly always nearly always contains more than one mode (“branched”), respectively. Only within a narrower range of intermediate fecundities does branching undergo gains and losses over short enough time scales to be biologically relevant (Figure 3).

Our framework allows us to confirm that the trait mean in the population can spend time away from its equilibrium value—the potential branching point due to drift, even when selection is stabilizing (see figure 1(c);(f)). Claessen et al. (2007) mentioned this phenomenon as one of the reasons for the delay or even prevention of branching in small populations. This fact, however, does not enter in our analysis of the potential for branching: we only use the equilibrium value of the trait mean, 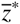, when calculating the expected change in the variance *μ_v_* and the stationary distribution of the variance *p_v_*. Furthermore, for the parameters investigated in figure 3, we overestimate the minimal population size at which branching reliably occurs relative to simulations, which suggests that the wandering of the trait mean 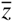 near 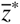 may not strongly prevent branching. It may, however, affect the timing of branching, as suggested by Claessen et al. (2007), a point that we do not investigate.

Assuming that both population size *N* and the trait mean 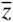 are at equilibrium is not the only approximation made to determine whether evolutionary diversification can occur or not. We also assume that the distribution of traits is Gaussian—which is clearly not the case after branching, if it happens. In addition, while we use the fact that the trait is bounded between 0 and 1, these boundaries are actually not reflected in the mutation kernel, which assumes that mutational effects have the same mean and variance regardless of the trait of the parent. Despite these limitations, our approximation of the stationary distribution of the variance allows us to identify the three branching regimes observed in the simulations: when drift prevents branching, when branching occurs intermittently, and when branching is virtually certain.

To conclude, our study confirms that evolutionary diversification can occur in asexual populations of finite size (although not in a randomly mating sexual population, see Appendix E). Even though random drift, which is stronger in smaller populations, may hinder diversification, evolutionary branching may nevertheless occur provided that selection is diversifying and strong enough to counter drift or if enough variation is introduced by mutation. Even if branching can occur, however, our results indicate that populations of intermediate size will tend to be branched only for transient periods of time, undergoing repeated cycles of branching and collapse. Thus, diversifying selection must be strong enough relative to drift, not only to allow evolutionary branching but also to allow branched populations to persist long enough to contribute substantially to the speciation process.

## 5. Acknowledgements

We thank the Otto lab members for feedback and L. Lehmann for discussions. Funding was provided by a Natural Sciences and Engineering Research Council Discovery grant (to SPO), CREATE Training Program in Biodiversity, a fellowship of the Wissenschaftskolleg zu Berlin and an Agence Nationale de la Recherche grant ANR-14-ACHN-0003 (to FD). We are grateful to the INRA MIGALE bioinformatics platform (http://migale.jouy.inra.fr) for providing computational resources.

## Appendices

## A. Derivation of the diffusion terms

In this section, we detail the calculations of the expected changes and squared changes in population size (*N*), trait mean 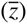 and variance (*v*), the latter two being defined as

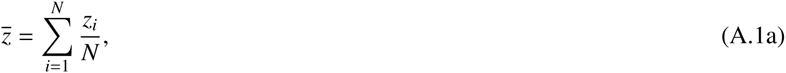

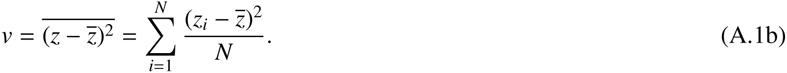

### A.1. Realised changes

Suppose that there are *N* individuals in the population at time *t*, and that the vector **z** = *{z*_1_*,···, z_N_*} contains the states of all individuals at time *t* (their order does not matter).

#### A.1.1. Birth

Let us assume that during a small time interval, individual *i*, with trait *z_i_*, reproduced and produced an offspring with trait *z_i_* + *δ*. Then at the next time step, the size of the population is *N* + 1, and the state of the population is represented by a vector ***z′*** of size *N* + 1, **z′** = {**z**, *z_i_* + *δ*}. In this case, the changes in population size, trait mean, and trait variance are as follows:

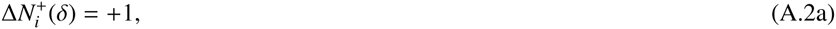

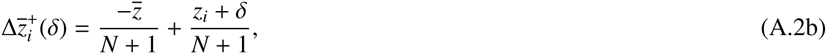

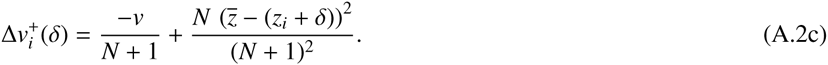

(Note that (A.2c) is obtained after a few lines of calculations.)

#### A.1.2. Death

If the event was instead a death, and individual *i* dies, the size of the population at the next time step is *N* – 1, and the corresponding changes are

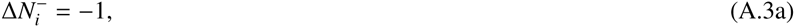

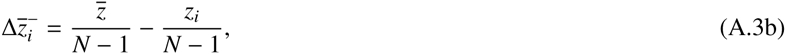

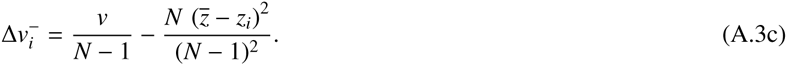

### A.2. Probability of a given change

We denote the death propensity of an individual *i* by *D_i_* and its propensity to reproduce (i.e., its fecundity) by *F_i_*. 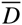 and 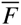 give the mean values of these propensities within the population.

#### A.2.1. Birth

The probability that the next event is a birth is

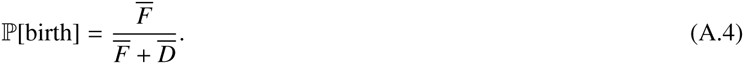

Given that the next event is a birth, the chance that individual *i* reproduces is

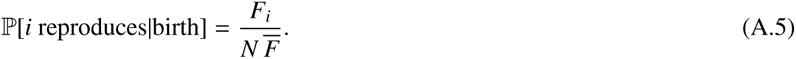

#### A.2.2. Mutation

The probability that the difference between *i*’s trait and its offspring’s trait is *δ* is given by *u*(*δ*), the mutation kernel. We assume that mutation is unbiased: the expected deviation is 0 and we denote by 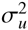 the mutational variance:

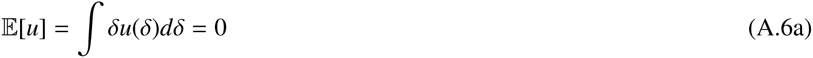

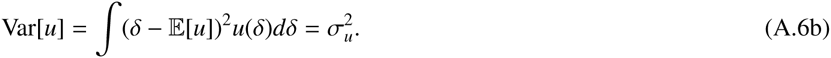

#### A.2.3. Death

The probability that the next event is a death is

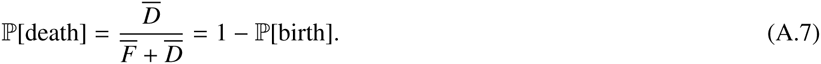

We assume that survival is density-dependent, but type independent. In a population of size *d*, the death propensity of every individual is a linearly increasing function of the population size with slope *d*:

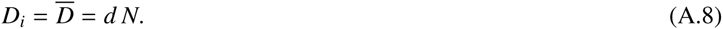

Given that the next event is a death, the chance that individual *i* dies is

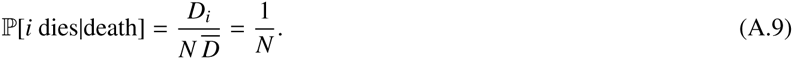

### A.3. Assumptions on the distributions of traits

In general, an individual’s fecundity will be a function of its own trait, *z_●_*, and of the distribution of traits in the population. We will assume that this distribution is Gaussian and of small variance. This means that knowing the trait mean 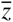 and variance *v* entirely characterizes the distribution of traits and that we can Taylor-expand individual fecundity as follows:

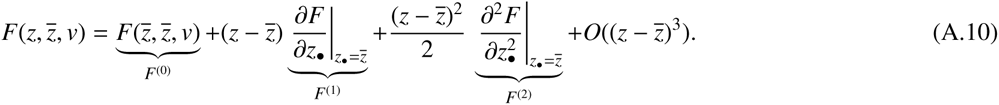

In particular, this means that we can write

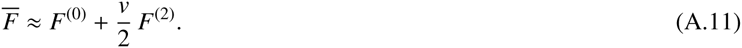

### A.4. Conditional changes

Conditioning on a state (*N,z*) of the population at time *t*, we denote by 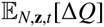 the expectation of a change in some quantity *Q*, between two time steps:

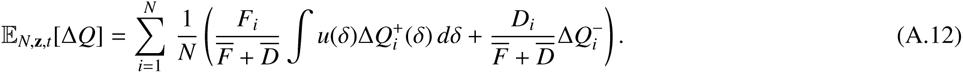

This formula uses the law of total probabilities. The values of 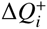 and 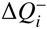 are given by (A.2) and (A.3) for *N*, 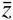, and *v*.

#### A.4.1. Population size

Substituting (A.2a) and (A.3a) into (A.12) and simplifying, we obtain

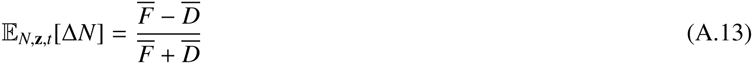

for the expected change. We also find

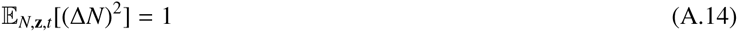

for the expected squared change.

We can further simplify equation (A.13) using the definition of *D_i_* given in (A.8) and the approximation of 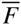 for small variance given in (A.11):

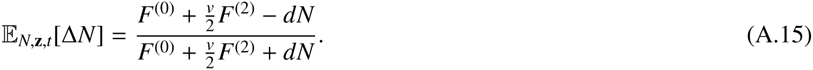

Equation (A.13) also implies that at demographic equilibrium, 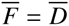. We will use this result in the following.

#### A.4.2. trait mean

*Expected change:*. Substituting (A.2b) and (A.3b) into (A.12) and simplifying, we obtain

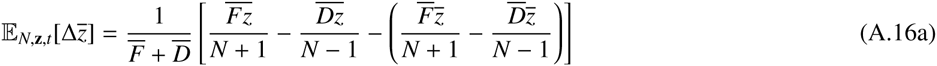

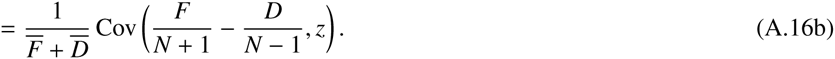

This is a version of the Price equation (Price, 1970; Frank, 2012). This gets clearer if we write *w_i_*, the expected proportion of the population, at the next time step, corresponding to an individual *i* and its potential offspring:

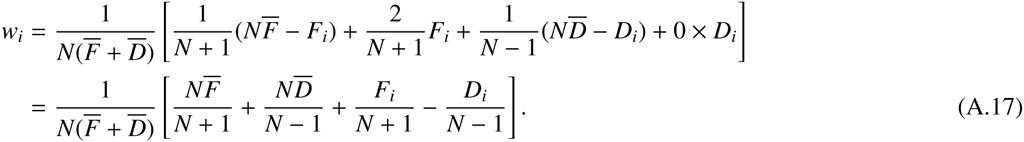

We note that (A.17) implies that

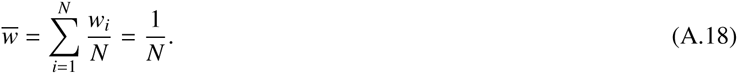

Using (A.17) and (A.18), we can rewrite (A.16b) in the familiar Price equation form (Price, 1970; Frank, 2012):

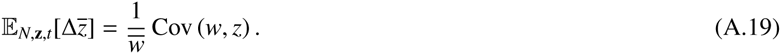

At demographic equilibrium, 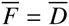, and we can rewrite (A.16b) as follows:

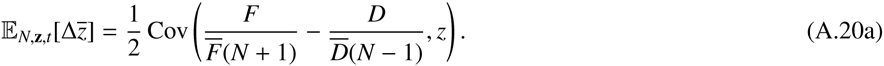

Using (A.8) and (A.11), this reduces to

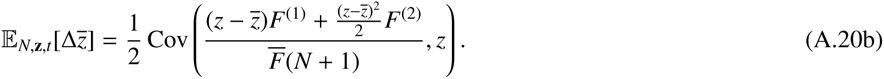

Finally, using the fact that the distribution of traits is assumed to be Gaussian, so that 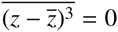, we obtain

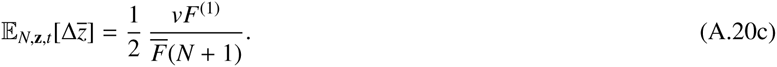

*Expected squared change:*. The expected squared change in the trait mean is

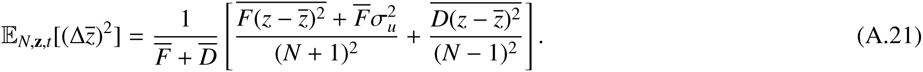

At demographic equilibrium, this simplifies to

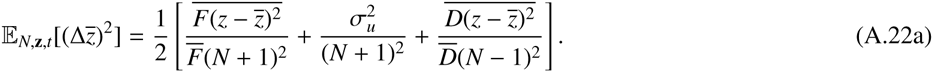

Using (A.8) and (A.11) and the fact that the distribution is assumed to be Gaussian (so that 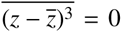 and 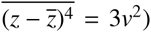, this reduces to

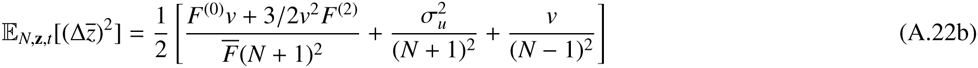

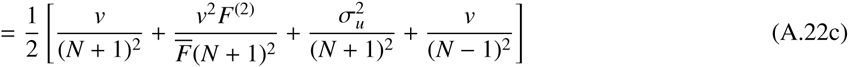

and finally, neglecting the term in *v*^2^,

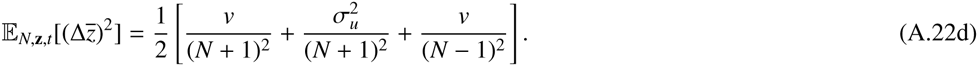

#### A.4.3. Trait variance

*Expected change:*. Substituting (A.2c) and (A.3c) into (A.12) and simplifying, we obtain

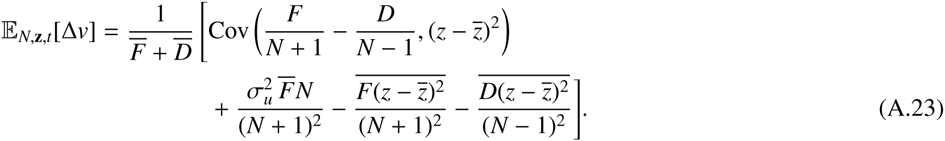

The first line corresponds to what is describes as the effect of “selection” in the Price equation, while the second line corresponds to “transmission”. We note that the “transmission” line also contains terms that do not correspond to mutation because the squared distance to the mean changes even in the absence of mutation (because the mean itself changes).

At demographic equilibrium, where 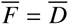, (A.23) reduces to

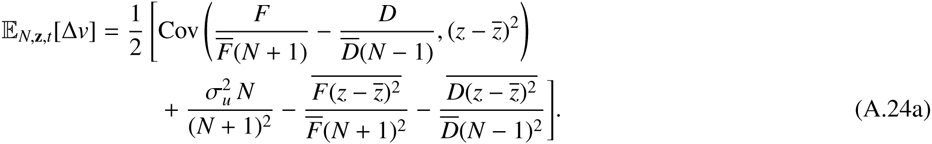

Using (A.8) and (A.11),

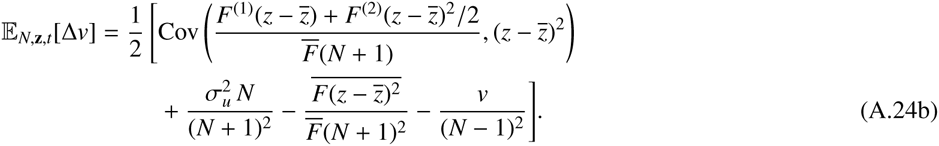

Now using the fact that the distribution is Gaussian:

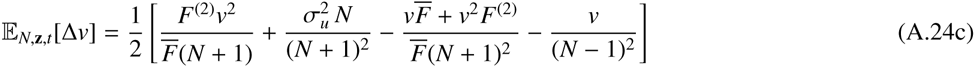

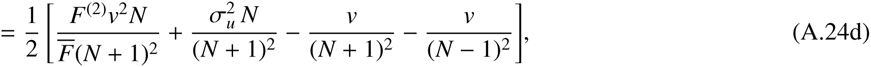

which leads to equation (11) in the main text. When population size *N* is large and trait variance *v* is small, this can be rewritten as

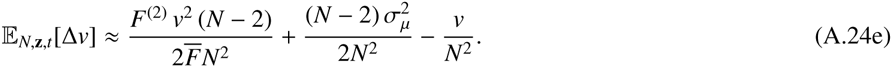

**In the absence of selection** In the neutral case, *F*^(2)^ = 0, and we are left with

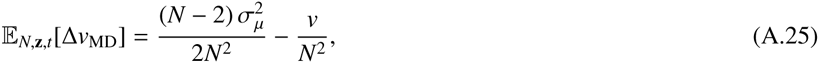

where MD stands for mutation-drift, the two evolutionary forces left. We denote by 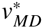 the variance at the mutation-drift equilibrium, such that the above equation is equal to zero:

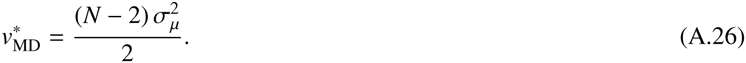

*Expected squared change:*. We now consider the expected squared change in the variance:

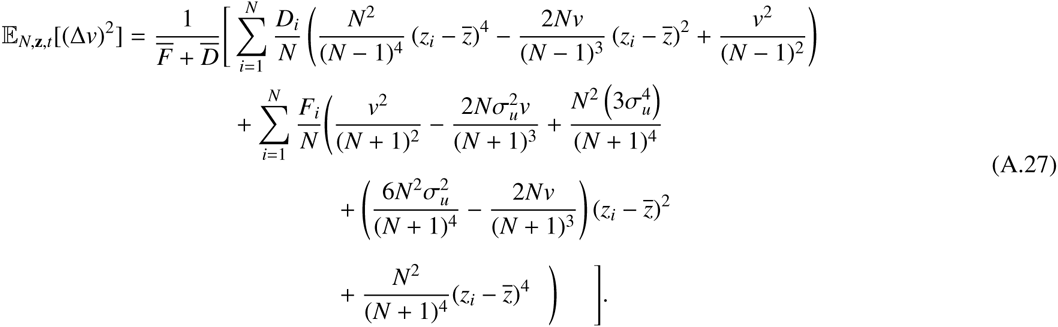

We then simplify the *F_i_* and *D_i_* terms using (A.8) and (A.11), and we use the assumption of a Gaussian distribution of traits, noting that in this case 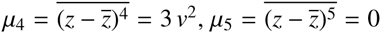 and 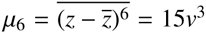. At demographic equilibrium, equation (A.27) becomes

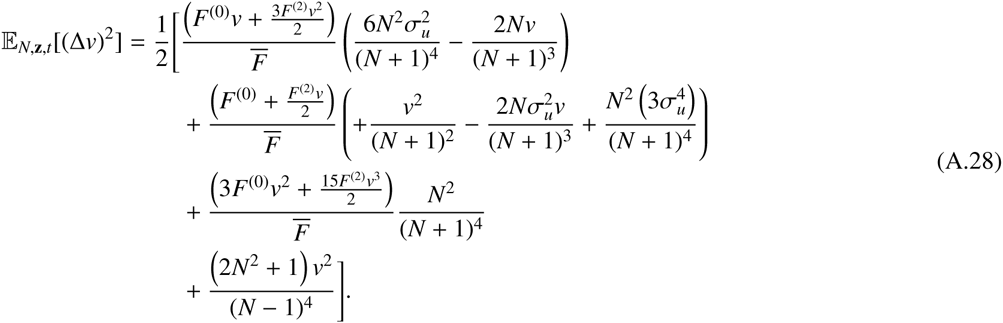

This expression can be further simplified by neglecting terms of order *v*^3^ and higher, *v*^2^*σ_u_* and higher, and 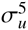 and higher. Also, we can approximate the expression for relatively large population sizes (small 1/*N*), and we obtain

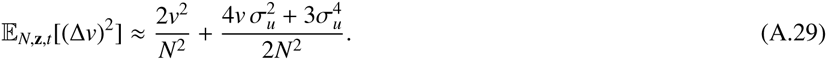

It turns out that the shape of the stationary distribution of the variance will almost not be affected if we use an even simpler expression, neglecting the effect of mutation on the expected change in the trait variance:

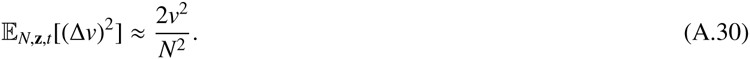

## B. Social evolution example

We illustrate our results with a specific example, a model of social evolution in a well-mixed population. In this example, an individual’s fecundity depends on both its type (*z*) and the distribution of types in the population (through its mean 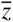 and variance *v*) and is given by

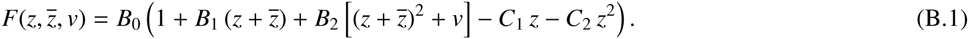

The *F*^(^*^k^*^)^ terms, defined in (A.10), are as follows:

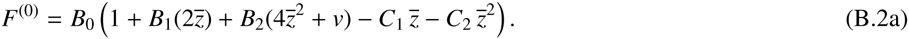

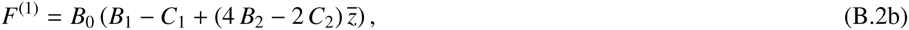

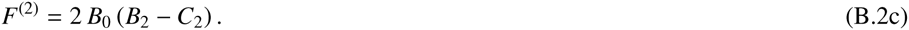

The expressions of the stationary distributions are obtained by integrating equation (2) for each variable, as described in the main text. These integrations are done with *Mathematica*. Although explicit solutions are found, their expressions are too complicated to present here, but they are available in a Supplementary *Mathematica* file.

## C. Evolutionary stability and convergence stability in Wakano and Lehmann (2012)

In Wakano and Lehmann (2012), invasion conditions are calculated using the stationary average frequency of the (mutant) allele *z_M_* that appeared in a population fixed for the allele *z_R_*, denoted by *ρ*(*z_M_*, *z_R_*). When the mutation is neutral, *ρ*(*z_M_*, *z_R_*) = 1/2; otherwise, the allele *z_M_* is said to successfully invade when its stationary average frequency is greater than the stationary average frequency of a neutral mutant, 1/2. The quantity *ρ*(*z_M_*, *z_R_*) − 1/2 is then used as if it corresponded to invasion fitness in order assess convergence and branching properties.

The selection gradient at a strategy *z* is defined as

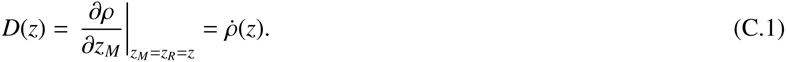

A singular strategy *z*^*^ is a strategy at which the selection gradient vanishes:

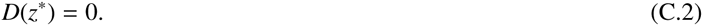

The convergence stability condition in Wakano and Lehmann (2012) is *Q_CS_* < 0, with:

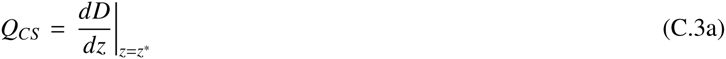

Using the formula of total derivatives, we can rewrite this as

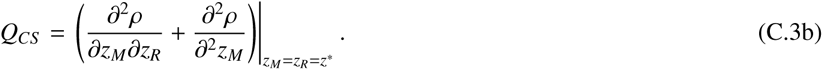

The evolutionary stability condition in Wakano and Lehmann (2012) is *Q_TS_* < 0, with:

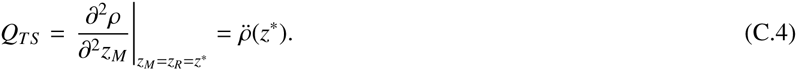

The relationship between *Q_TS_* and *Q_TS_* can be simplified by using the fact that *ρ* is a stationary average frequency and that we are considering two alleles:

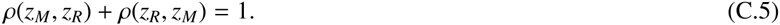

Equation (C.5) implies that (taking the derivative with respect to the first argument):

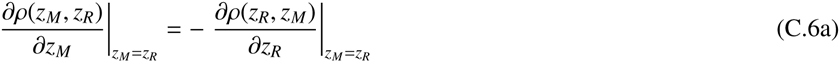

and then that (taking the derivative with respect to the second argument)

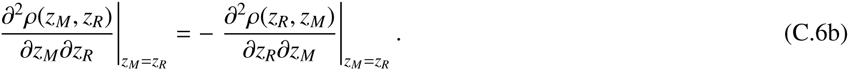

Because the partial derivatives commute, and because we are evaluating the derivatives when *z_M_* = *z_R_*, then equation (C.6b) implies that

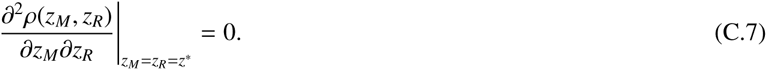

Given the definitions of *Q_TS_* (equation (C.4)) and *Q_CS_* (equation (C.3b)), the result shown in (C.7) means that for any life-cycle and any population size

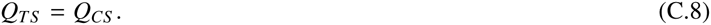

In other words, when using the stationary average allele frequencies to describe invasion fitness, the conditions for evolutionary stability and convergence stability must match, which indicates that this surrogate for invasion fitness is unable to identify the conditions under which evolutionary branching occur (i.e., where *Q_CS_* < 0 while *Q_TS_* > 0).

## D. Oligomorphic dynamics

### D.1. Model

In this section, we present an analysis of a deterministic version of the model, using oligomorphic dynamics (Sasaki and Dieckmann, 2011; Débarre et al., 2013), which allows us to determine when a trait distribution will be multimodal. Under this approximation, the distribution of traits in the population is decomposed into a sum of unimodal distributions, each corresponding to a “morph” – this approximation extends the adaptive dynamics framework to incorporate standing genetic variance within each morph.

There are *M* morphs in the population (technically, in our case it will be sufficient to consider *M* = 1 and *M* = 2 only). The density at time *t* of individuals belonging to morph *m*, having trait *z* is denoted by *ϕ_m_*(*z, t*). Its dynamics are described by the following equation

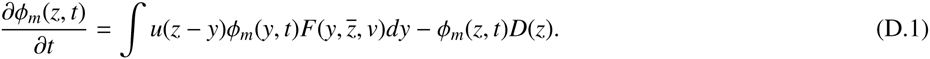

Notation is similar to in our main model: *u* is the mutation function (see section A.2.2), *F* is the fecundity function, and *D*(*z*) = *dN* represents density-dependent mortality. As previously, we drop the time dependence to simplify the notation.

*Dynamics of the total population size:*. Using equation (D.1), we can write the dynamics of *N* as follows:

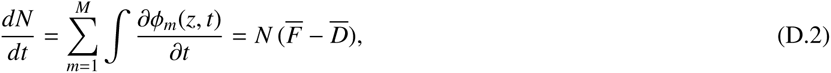

an equation similar to equation (A.13) (the difference is a factor 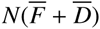, scaling the time between events). At demographic equilibrium (*dN/dt* = 0), we have 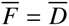.

*Dynamics of the size of morph m:*. The dynamics of total size of morph *m* are as follows:

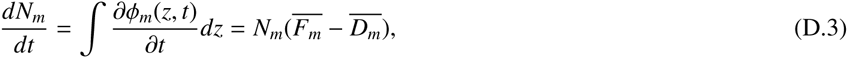

where 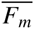 and 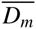 are the mean fecundity and mean mortality within morph *m*.

*Dynamics of the relative size of morph m:*. The proportion of the population consisting of morph *m* is *q_m_* = *N_m_/N*, and its dynamics are as follows:

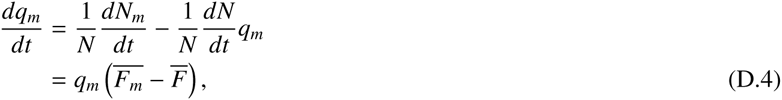

because mortality is type- and frequency-independent, so that 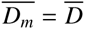.

*Dynamics of the frequency of a trait within morph m:*. We denote by *p_m_*(*z, t*) the frequency of individual with trait *z* within morph *m*. Its dynamics are as follows:

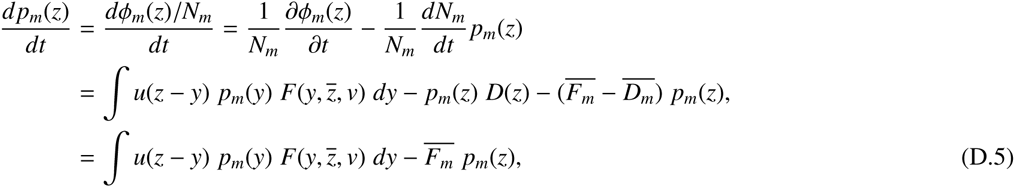

again using the fact that mortality is the same for all individuals in the population at a given time.

### D.2. Dynamics of the moments

We can now use equation (D.5) to compute the changes in the trait mean and trait variance within each morph. For this, we will use the same kind of second order expansion of the fecundity function as in equation (A.10), except that the approximation is now done for each morph, near the morph’s trait mean:

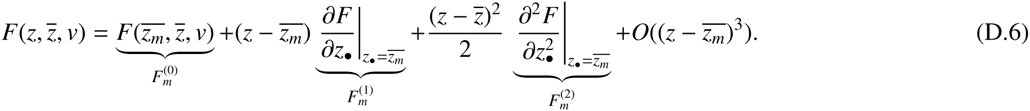

This means that we assume that the variance *v_m_* within morph *m* is small – but this is not necessarily the case for the overall variance *v* in the population. With this approximation, we can rewrite the mean fecundity within morph *m* as follows:

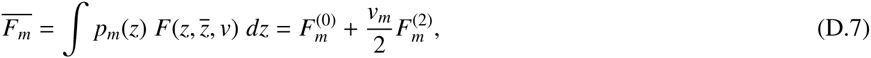

where *v_m_* is the variance within morph *m*.

#### D.2.1. Changes in the trait mean within morph m

We have

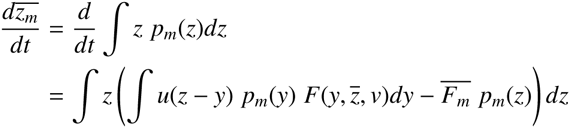

We now use the expansion of equation (D.6), and we obtain:

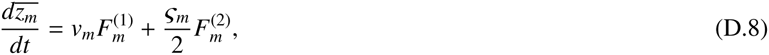

where *ζ_m_* is the third central moment of the distribution of traits within morph *m*. We can now assume that the distribution of traits within each morph is Gaussian, so that *ζ_m_* = 0, and we end up with the within morph, deterministic, equivalent of equation (A.20c):

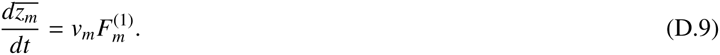

#### D.2.2. Changes in the variance within morph m

We have

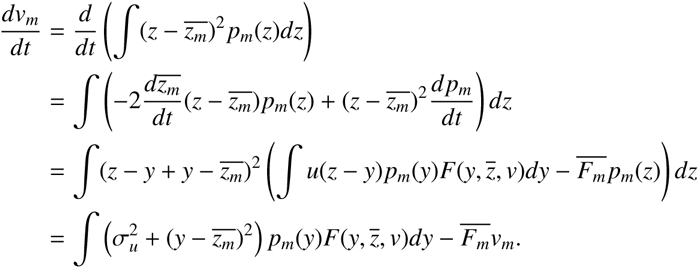

Now using the expansion of equation (D.6), we obtain

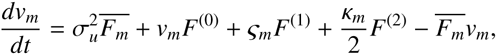

where 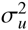 is the mutational variance (see equation (A.6)), and *κ_m_* the fourth central moment of the distribution of traits within morph *m*. As previously, we assume that the distributions within each morph are Gaussian; consequently, *κ_m_* = 3*v_m_* (and *ζ_m_* = 0), and we obtain.

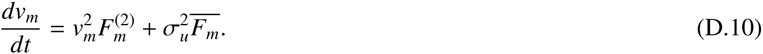

The first term in equation (D.10) corresponds to selection, via the curvature of the fecundity function, and the second term corresponds to the variance contributed by mutation. Because we are here considering a population of infinite size, there is no effect of drift.

As previously, we note that the dynamics of the population size is of order 1 (equation (D.3)), the dynamics of the trait mean within morph *m* is of order *v_m_* (assumed to be small), and the dynamics of the variance within morph *m* of order 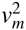: we can hence again decompose time scales, assuming that population size equilibrates first, followed by the trait mean, followed by the trait variance.

### D.3. Dynamics with one morph

Let us start with one morph in the population (*M* = 1); we can drop the *m* subscripts here. As population size equilibrates, 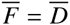 (equation (D.3)). Then, the trait mean of the morph equilibrates, which occurs at a value *z*^*^ for which *F*^(1)^ = 0 (equation (D.9)). Whether selection is then stabilizing or diversifying depends on the sign of *F*^(2)^ at *z*^*^ (equation (D.10)).

### D.4. Dynamics with two morphs

Let us consider the case of diversifying selection (*F*^(2)^ > 0), and let us now represent the distribution of the trait as the sum of two Gaussian distributions (indexed by *m* ∊ {1, 2}), currently having the same mean *z*^*^ and the same variance *v_m_*. Densities within each morph equilibrate fast, and for each morph, 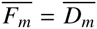. The trait means change according to equation (D.9). Variances within a morph change much more slowly, so we can first approximate them as constant. Note that this does not assume that the global variance *v* changes at the same rate, because the global variance will be driven by changes in the trait means of each morph. (Throughout, we discuss the global distribution of traits as the sum of the distributions for each morph.)

### D.5. Social evolution example

#### D.5.1. With one morph

The fecundity function and its derivative are the same as in Appendix B, equation (B.2). The singular strategy *z*^*^ is found by solving *F*^(1)^ = 0, which yields the same value as in the main text (equation (10)). This value is an attractor when 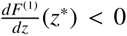, which occurs when 2*B*_2_ – *C*_2_ < 0. Selection is diversifying, i.e., the variance of the distribution can increase due to selection when *F*^(2)^(*z*^*^) > 0, which occurs when *B*_2_ – *C*_2_ > 0.

#### D.5.2. With two morphs

We can rewrite an individual’s fecundity function by decomposing the population into morphs; equation (B.1) then becomes

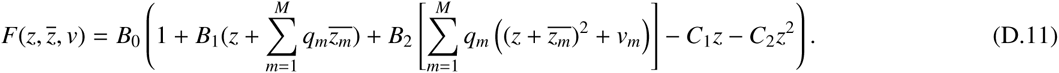

As a reminder, *q_m_* = *N_m_/N*: it is the relative density of morph *m* in the population, and 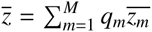. We now consider the case where there are two morphs in the population, *M* = 2.

Then, the first derivative of *F* with respect to the trait of the focal individual, evaluated at 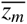, reads

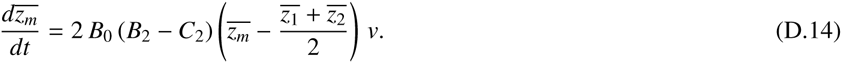

We can now rewrite equation (D.9), evaluating *N* (total population size) and the *q_m_* terms (proportion of morph *m* in the population) at equilibrium, since densities equilibrate fast. The demographic equilibrium with the two morphs is calculated by solving *dN/dt* = 0 for *N* (equation (D.2)) and, for *m* = 1 and *m* = 2, by solving *dq_m_/dt* = 0 (equation (D.4)) for *q_m_*. To simplify the calculations, we also assume that the variances of the two morphs are equal: *v*_1_ = *v*_2_ = *v*. We find that at demographic equilibrium, the proportion of the population belonging to morph 1 is given by

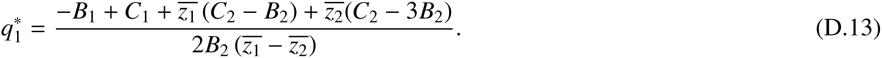

We note that when the two morphs are equidistant from the one-morph equilibrium value *z*^*^, equation (D.13) simplifies to 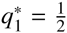. We finally obtain

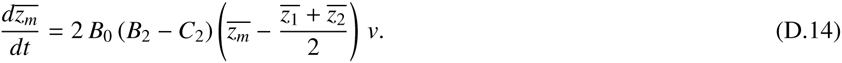

We are first considering the initial case where the means of the two morphs are both at *z*^*^ (i.e., the population has not started diversifying yet and is still unimodal); in this case the two means evolve away from *z*^*^ if *B*_2_ – *C*_2_ > 0, which is precisely the condition for selection to be diversifying at *z*^*^ (see equation (B.2c)).

The only equilibrium of Equation (D.14) occurs when 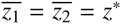, that is, when there is actually only one morph. That is, we cannot identify the equilibrium to which the system heads when selection is diversifying. This is because we have not explicitly taken into account the fact that the trait is bounded (0 ≤ *z* ≤ 1). What actually happens is that the two morph means keep evolving away from *z*^*^, until they reach the boundaries 0 and 1, the extremum values of the trait.

## E. Sexual reproduction

### E.1. Assumptions

The model presented in the main text assumes clonal reproduction: only one parent is required for reproduction. The framework can however be extended to accommodate sexual reproduction. For this, we need a map linking the parent’s phenotype to their offspring’s phenotype. Here, we will assume that an offspring born to parents with phenotypes *z_i_* and *z_j_* has a phenotype *z_o_* given by

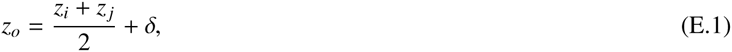

where *δ* is a random variable with mean 0 and variance 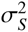, which takes into account the effects of mutation, segregation and recombination.

All individuals in the population are assumed to be hermaphoditic. As previously, an individual with trait *z_i_* is chosen to reproduce with probability given by 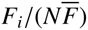. This individual now has to choose a mate. We assume that mating occurs at random, and self-fertilization is possible: a mating individual *i* chooses individual *j* with probability 1/*N*, *N* being the current size of the population. We can hence write the expected value of the offspring phenotype of a reproducing individual *i*, *E_i_*[*z_o_*] (and also of its square, 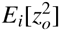, integrating over all possible mates and all possible values of *δ*:

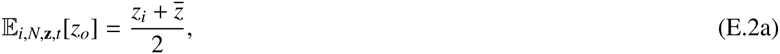

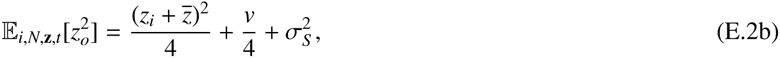

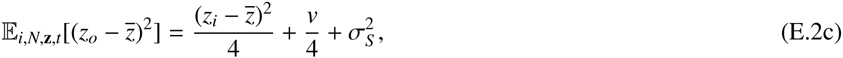

where 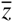 and *v* are the mean and variance of the distribution of traits in the population.

### E.2. Expected changes

#### E.2.1. Population size

The mode of reproduction does not change the expected changes in the size of the population: equation (A.15) remains unchanged. As previously, we consider the case where the population has reached its demographic equilibrium, meaning that 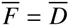.

#### E.2.2. Trait mean

The mode of reproduction however affects the expected changes in the moments of the distribution of traits. Using equation (E.2a), the change in the trait mean is as follows:

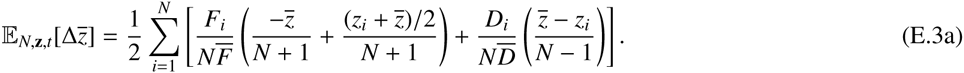

Using the expansion from equation (A.10), this further simplifies into

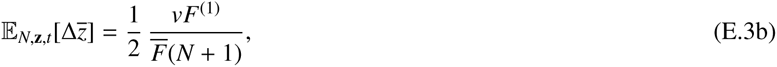

an expression that differs from its clonal equivalent (equation (A.20c)) by only a factor 1/2.

#### E.2.3. Trait variance

Using equation (E.2c), we can write the expected change in the trait variance as follows:

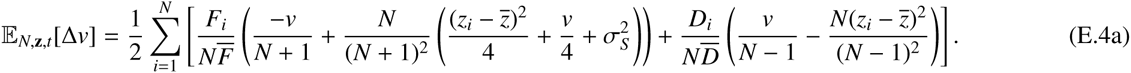

Using the expansion from equation (A.10), this further simplifies into

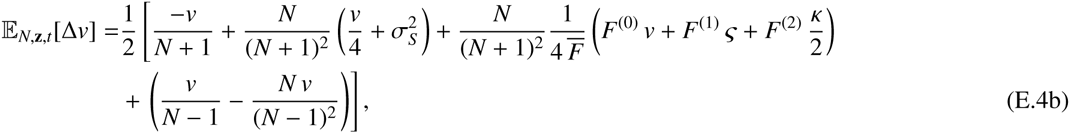

where *ζ* and *κ* are the third and fourth central moments of the distribution of traits, respectively. Assuming as previously that this distribution is Gaussian, we have *ς* = 0 and *κ* = 3*v*^2^, and we obtain after simplification:

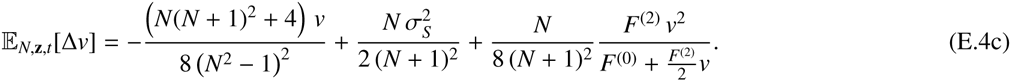

Identifying the first term of equation (E.4c) as the drift term, we can now compare it to its clonal equivalent (equations (A.24d) or (11) in the main text). In the clonal model, the drift term is of order 1/*N*^2^, while the mutation and selection terms are of order 1/*N*; thus, the drift term becomes relatively unimportant when the population size becomes large. The situation is different in the sexual model (equation (E.4c)): the “drift” term (which now also includes the randomness of choosing a mate) is of order 1/*N*, the same as the mutation and selection terms. Thus, diversifying selection (reflected in the last term of (E.4c)) will only have a modest effect on the mutation-recombination-drift balance obtained by setting the first two terms to zero, and it will not be possible for *v* to increase by orders of magnitude from the initially small values. As a result, the drift term does not vanish when population size becomes large. Consequently, evolutionary diversification is precluded, as has been found previously from deterministic models of a single population with random mating (Dieckmann and Doebeli, 1999).

## Supplementary figure

**Figure S1:**
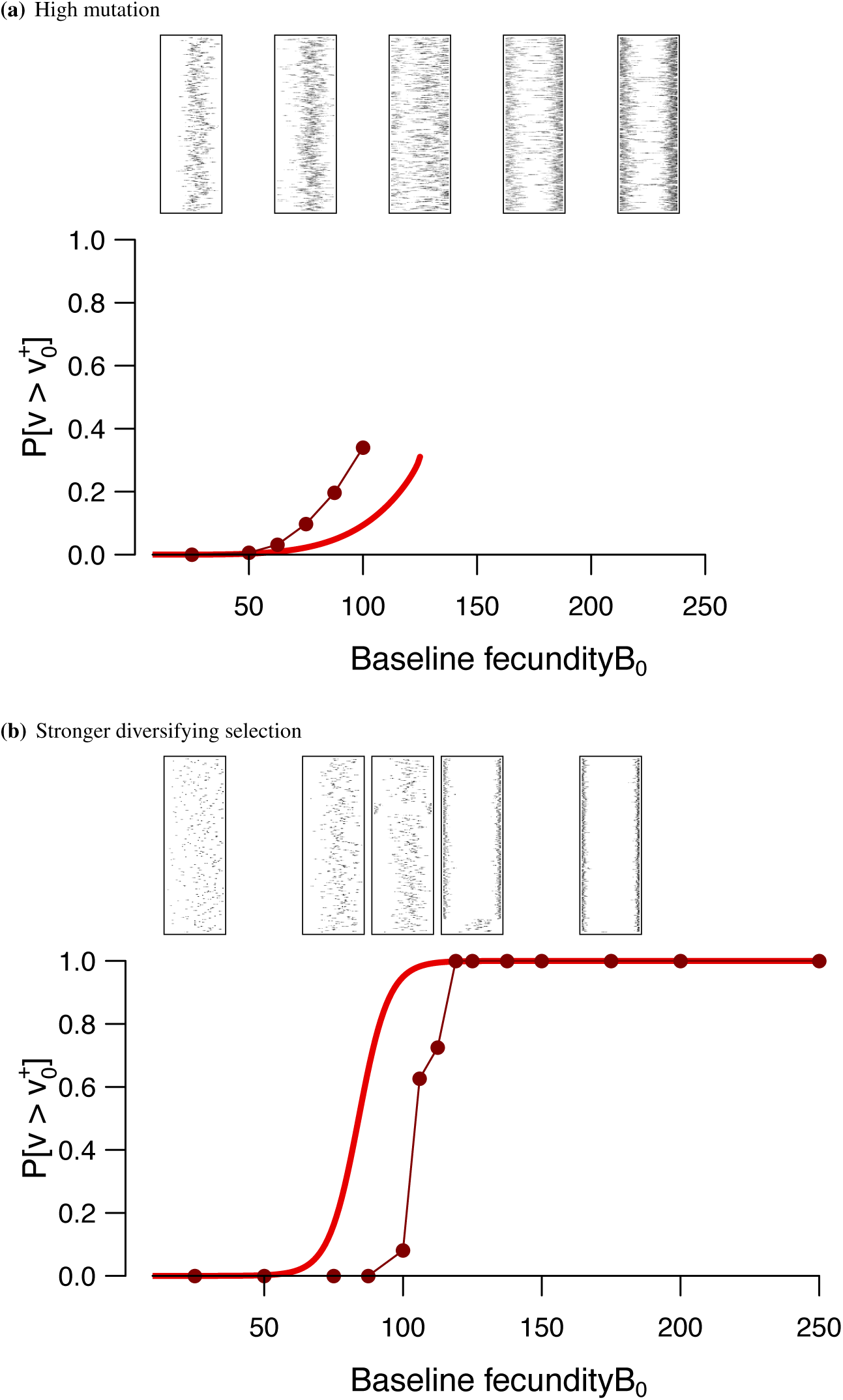
Equivalents of figure 3, with different parameters. In (a), the mutation probability is much higher:*μ*_0_ = 0.1 (instead of 0.01), so that σ*_μ_* ≈ 0.00632. There are no data points after *B*_0_ = 125, because value corresponds to (branching becomes certain). In (b), diversifying selection is stronger (*B*_2_ = −0.9, *C*_2_ = −1.6, *B*_1_ = 4.8, *C*_1_ = 4.56).

## Footnotes

1 Provisional DOI: doi: 10.5061/dryard. b3604

